# The Effect of Refractive Blur in the Vividness of Mental Imagery

**DOI:** 10.1101/2020.07.17.208017

**Authors:** Tharan Suresh, Anupama Roy, Atif Shaikh, Joanne Lydia Rajkumar, Vivek Mathew, A.T. Prabhakar

## Abstract

**Background:** Visual mental imagery or “seeing with the mind’s eye” is an everyday phenomenon. Visual mental imagery and visual perception share common neural-networks. Hence deficits that affect the visual perception may also affect visual mental imagery.

**Aim:** We aimed to study the effect of refractive blur on the vividness of mental imagery.

**Methods:** Subjects were recruited from volunteers and divided into two groups; individuals with refractive errors–Ametropes(AM), and individuals without refractive errors – Emmetropes(EM). After filling in the Verbalizer-Visualizer-Questionnaire (VVQ), the subjects were asked to perform a mental imagery task with and without refractive blur. The participants were asked to generate a mental image of a specific object initially with eyes closed, eyes open and then with refractive blur in random order, and then judge the vividness of the mental image on a Likert scale ranging from 1 (low vividness) to 5 (good vividness). The EM participants had to wear a + 2D spectacles to produce refractive blur.

**Results:** A total of 162 participants were recruited to the study. Of these 73 were EM and 89 were AM. Of the AM, 30 had additional astigmatism. The mean VVQ score was 64.9(11.2). The mean refractive error was 1.8(1.3)D. Following the mental imagery task, at baseline with eyes closed, 138 (85.5%)subjects had vivid mental imagery close to visual perception(Likert scale:5). With the opening of the eyes, the vividness dropped by at least 1 point in the Likert scale in 139(85.8%). With the introduction of refractive blur, 153(94.4%) subjects had a drop in the vividness of the image by at least 1 point and 22(13.6%) subjects by at least 2 points.

**Conclusion:** Introduction of refractory blur results in the reduction of the vividness of mental imagery.

## Introduction

Imagination is one of the prime higher-order cognitive functions in everyday life and visual mental imagery forms one of its key component(Pearson et al., 2011). We use visual mental imagery in our day-to-day activities to accomplish various tasks. This includes memory, learning, navigation, decision making analytical thinking, creativity and art (Keogh & Pearson, 2011; Paivio, 1969).

The phenomenon of imagination or ‘seeing with the mind’ can be explained as internally generating an image (visual representation) without an external visual stimulus(S. Kosslyn, 1987). The similarities of this phenomenon to visual perception has been vastly studied over a long period by philosophers and neuroscientists leading to a debatable conclusion that both visual perception and mental imagery share common neural pathways(S. M. Kosslyn, 1994). We hypothesised that, if there is a common neural substrate for mental imagery and visual perception, a perceptual defect affecting visual perception would increase the load on the common neuronal networks, and thus may affect mental imagery. A refractive error causes impairment in the clear perception of visual representation resulting in a visual perceptual deficit. We aimed to study, whether an introduction of a refractive blur affects the vividness of the in the internally generated mental image.

## Materials and Methods

### Subjects

We recruited healthy volunteers after informed consent. The study was conducted in accordance with the Declaration of Helsinki. The participants had no neurological or psychiatric disorders and were blinded to the purpose of the experiment. The eyesight of every subject was examined at a tertiary care ophthalmology unit. The subjects were examined for visual acuity; a certified ophthalmologist assessed refraction, visual field, retinal examination and colour vision. Based on the ophthalmology results, the subjects were divided into two groups – Ametropes (subjects with refractive error) and Emmetropes (subjects without refractive error –normal vision) similar to the study by Palmero et al(Palermo et al., 2013).

### Verbaliser-Visualiser Questionnaire (VVQ)

Verbalizer-Visualizer Questionnaire (VVQ) is a self-administered questionnaire consisting of 15 true-false items(Antonietti & Giorgetti, 1998). After the eyesight examination, the subjects were asked to fill the Verbaliser-Visualiser Questionnaire (VVQ). The VVQ assessed the subjects’ mental imagery function for various modalities like form, colour, shape, detail and spatial orientation. A score of 5 corresponded to a clear and vivid mental image, and a score of 0 corresponding to a very unclear/blurred image or an unimaginable scenario. The total score of the VVQ was calculated.

### Introduction of refractive blur

Immediately after the scoring was completed, the subjects were asked to perform a mental imagery task with and without refractory blur. The participants were asked to generate amental image of a specific object initially with eyes closed, eyes open and then with refractive blur in random order and then to judge the vividness of his mental image on a Likert scale ranging from 1 (low vividness) to 5 (good vividness). The EM participant had to wear a + 2D spectacles to produce refractive blur. A time interval of 30 seconds to 1 minute was given between the trials. Apple, elephant, a red rose, and a house were some of the specific objects that the subjects had to imagine for the mental imagery task. **Fig.1** shows the graphical representation of the methodology with eyes open, eyes closed and with refractive blur.

**Figure. 1:**
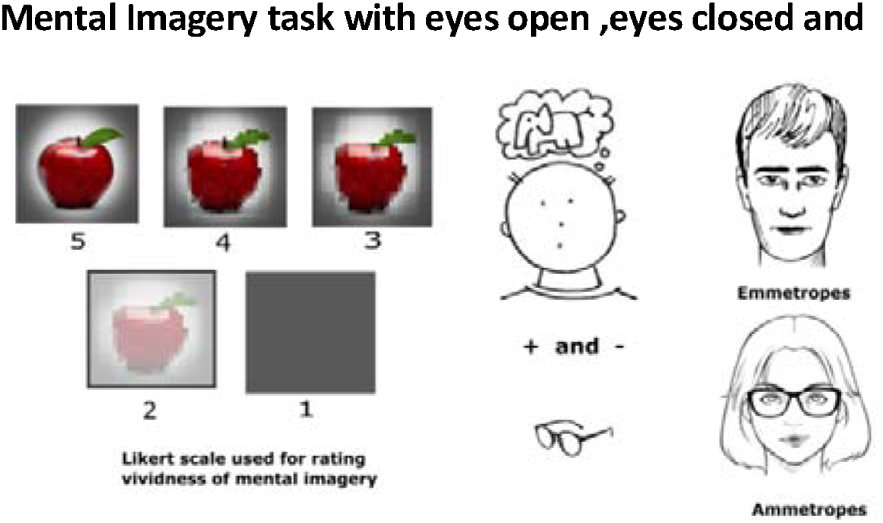
Ammetropes and Emmetropes were asked to perform a mental imagery task with eyes closed, eyes open and with refractive blur in random order, and the vividness of the mental imagery was recorded using the 5 point Likert scale. Refractive blur was introduced in the ammetropes by removing their corrective spectacles and in emmetropes by adding a +2 diopter corrective

The participants were comfortably seated in a well-lit and quiet room for filling up the VVQ with no time restrictions. The Imagery tasks were also performed in a similar room setting, allowing enough time for the subject to visualize in detail.

## Results

A total of 162 participants were recruited to the study. Of these 73 were EM and 89 were AM. Of the AM, 30 had additional astigmatism. The characteristics of the enrolled subjects is given in **Table 1**. The mean VVQ score was 64.9(11.2), 66.3(9.1) in EM and 62.5(14.3) in AM. The mean refractory error amongst the AM was 1.8(1.3)D. Following the mental imagery task, at baseline with eyes closed, 138 (85.5%)subjects had vivid mental imagery close to visual perception(Likert scale:5). With the opening of the eyes the vividness of the image dropped by at least 1 point in the Likert scale in 139(85.8%). With refractive blur, 153(94.4%) subjects had a drop in vividness of the image by at least 1 point and 22(13.6%) subjects by at 2 points**(Fig.2)**.

## Discussion

Mental imagery is conceived of as a type of top-down perception that functions like a weak form of perception(Lee et al., 2012). Behavioural data and functional brain imaging suggest that common sets of neural structures are employed during both processes(Pearson et al., 2011). Like visual perception, mental imagery seems to involve visual areas that are organized topographically(Kaas et al., 2010). The neural representation of mental imagery significantly overlaps with visual working memory and visual attention (Albers et al., 2013; Moriya, 2018). It has been previously demonstrated that visual working memory is limited both by visual information load and by the number of objects(Alvarez & Cavanagh, 2004). Behavioural data suggest that domains such as attention, recognition, and memory have capacity-limited buffers that can be viewed as competing map architectures(Franconeri et al., 2013). In this context, we postulated that if visual perception and mental imagery shared a common but limited neural substrate, introduction of a transient refractive blur should increase the information load and thus adversely affect the quality of the on-going mental imagery process. Vividness, that was used to assess the quality of mental imagery is defined as a construct expressing the self-rated degree of richness, amount of detail, and clarity of a mental image, as compared to the experience of actual seeing (D’Angiulli & Reeves, 2007). Vividness though a subjective measure, correlates with objectively quantifiable f-MRI activity in the visual cortex (Cui et al., 2007). Our results demonstrate that vividness of mental imagery was highest in the eye closed condition and dropped significantly with eyes open condition and further more after the introduction of refractive blur. This corroborates with our hypothesis that visual mental imagery and visual perception share a capacity-limited common neural substrate. Our results also show that the vividness of mental imagery as measured by the VVQ scores was higher in the EM as compared with the AM. This is similar to the results of Palermo et al and demonstrates that peripheral visual defects can affect mental imagery(Palermo et al., 2013). Our results correlate with the common experience of difficulty in imagining while multitasking. This could be explained by stating that generating mental images requires attention, and divided or fragmented attention would cause difficulty in generating mental images as a whole(Pearson et al., 2015). Mental imagery is a proven method to enhance neuro-rehabilitation, and mental imagery techniques have known to minimize the recovery period. Our results suggest that correction of peripheral visual defects including refractive errors may help in improving mental imagery in the context of rehabilitation(Evans et al., 2006). The limitations of our study included the use of VVQ, which may not be a pure measure of vividness of mental imagery(Antonietti & Giorgetti, 1998) and using only subjective behavioural data, without any f-MRI or electrophysiological correlates.

**Table 1:** Characteristics of the subjects.

|  | Total | Emmetropes (normal) | Ametropes (Refractive error) | p value |
| --- | --- | --- | --- | --- |
| Total number | 162 | 73 | 89 |  |
| Age | 27(6.8) | 26.6(7.2) | 27.6(6.4) | 0.356 |
| Gender (Male/female) | 84/78 | 38/35 | 46/43 | 0.220 |
| Refractive error |  | + 2 D | 1.8(0.3) | 0.268 |
| VVQ score | 64.9(11.2) | 66.3(9.1) | 62.5(14.3) | 0.042 |

**Figure 2:**
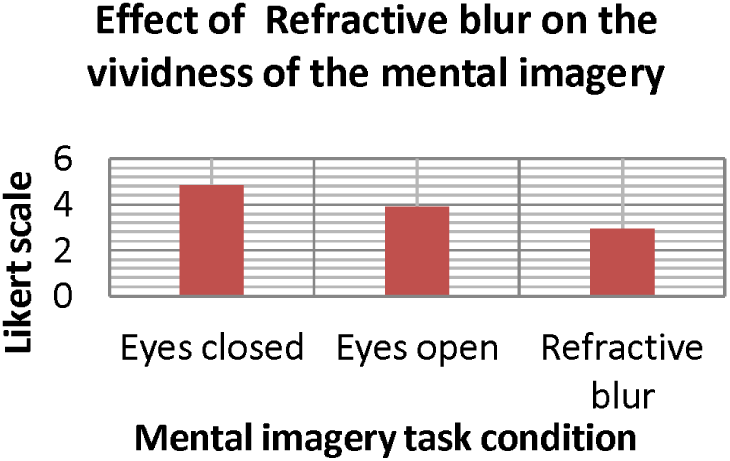
Ammetropes and Emmetropes were asked to perform a mental imagery task with eyes closed, eyes open and with refractive blur in random order, and the vividness of the mental imagery was recorded using the 5 point Likert scale. On introduction of a refractive blur, the vividness of mental imagery dropped as measured by the Likert scale.

**Table 2:** Results of the Mental Imagery Task. Kruskal Wallis test was performed on the data with the three conditions (Condition1: Mental imagery task with eyes open, Condition2: Mental imagery task with eyes closed and Condition3: Mental imagery task with refractive blur) as factors. The Kruskal-Wallis rank sum test showed a significant difference between the groups and effect size was calculated using epsilon squared (Kruskal-Wallis Chi-squared = 394.21, df = 2, p-value <0.0001, ⍰ ^2^= 0.813). Dunn Post-hoc test for multiple comparisons after Bonferroni correction showed significant differences between all three conditions with maximum significance between the eyes closed and refractive blur condition (Z=9.769, adj-p <0.001).

|  | Eyes closed | Eyes open | post refractive blur | P value |
| --- | --- | --- | --- | --- |
| Emmetropes (normal) | 4.87(0.36) | 3.9(0.27) | 2.93(0.53) | <0.00001 |
| Ametropes (Refractive error) | 4.83(0.4) | 3.84(0.36) | 2.94(0.44) | <0.00001 |
| Total | 4.8(0.4) | 3.8(0.31) | 2.93(0.49) | <0.00001 |

## Conclusion

In conclusion, our results show that when a refractive blur is introduced, the vividness of the internally generated image drops significantly thus supporting the hypothesis that visual perception and mental imagery share a common neural substrate.

## Acknowledgments

Nil

## Declarations

### Availability of data and material

Data will be shared on reasonable request

